# Persistent endotoxin exposure in macrophages elicits an immunometabolic profile susceptible to NF-κB and p53 perturbation

**DOI:** 10.1101/2025.04.30.651567

**Authors:** Chima V. Maduka, Xuan Xie, Ashley V. Makela, Evran Ural, Christopher H. Contag

## Abstract

The prolonged exposure of macrophages to endotoxin occurs in various chronic inflammatory conditions, including long-term inflammation triggered by particle-releasing implanted biomaterials, progressive pulmonary obstructive conditions, chronic bronchitis, colitis and hepatitis characterized by sustained exposure to gut-derived lipopolysaccharide (LPS). However, immune cellular metabolism in models of persistent inflammation remains undercharacterized, especially when compared to the well-studied immunometabolic changes following short-term LPS exposure. Here, we demonstrate that persistent LPS stimulation induces increased oxygen consumption alongside reduced levels of ATP and elevated glycolysis, with a mixed inflammatory profile indicative of endotoxin tolerance in primary bone marrow-derived macrophages. Pharmacological blockade of complex I of the mitochondrial electron transport chain lowered oxygen consumption and mitochondrial ROS by reducing NF-κB signaling. Surprisingly, NF-κB inhibition at TAK1 or IκBα phosphorylation, and p53 activation, reduced both oxygen consumption and glycolysis with minimal mitochondrial ROS impact. The absence of elevated mitochondrial membrane potential in prolonged LPS exposure suggests mitochondrial ROS production is likely reverse electron transport-independent, supported by the lack of effect of a mitochondrial depolarizer on oxygen consumption. These findings reveal a distinct immunometabolic phenotype in macrophage endotoxin tolerance, extending our understanding of immunobiology in chronic inflammatory diseases.

## Introduction

Macrophages are essential cellular mediators of inflammation, playing crucial roles in both tissue repair and regeneration^1^, as well as host defense against foreign agents. Under homeostatic conditions, macrophages primarily rely on mitochondrial oxidative phosphorylation (OXPHOS) for energy production. However, upon exposure to pro-inflammatory stimuli, macrophages rapidly change their energy dependence to glycolysis, a switch necessary for inducing a pro-inflammatory phenotype^2^. This process, termed metabolic (glycolytic) reprogramming, is triggered by the activation of macrophage pattern recognition receptors, such as Toll-like receptor 4 (TLR4), through interaction with pro-inflammatory agonists like lipopolysaccharide (LPS) from Gram-negative bacteria^3^.

Concomitant with elevated glycolytic flux, the tricarboxylic acid (TCA) cycle is disrupted^4^, accumulating inflammatory metabolites, especially succinate and citrate. Accumulated succinate stabilizes and activates the transcription factor hypoxia-inducible factor 1α (HIF-1α), which in turn drives the production of the pro-inflammatory cytokine IL-1β^5^. Simultaneously, citrate fuels the production of pro-inflammatory lipids, including leukotrienes and prostaglandins^4^, and mitochondrial function is redirected towards the generation of reactive oxygen species (ROS) at complex I of the electron transport chain through a process known as reverse electron transport (RET)^6-8^.

In resting macrophages, the transcription factor nuclear factor kappa B (NF-κB) is located in the cytoplasm as an inactive dimer (composed of the p65 and p50 subunits) bound to its inhibitor IκBα^9^. Macrophage activation triggers IκB kinase (IKK)-dependent phosphorylation and subsequent ubiquitination of IκBα, leading to the release and nuclear translocation of the active NF-κB dimer^10^. Within the nucleus, NF-κB regulates gene expression, promoting the production of pro-inflammatory cytokines and chemokines, including MCP-1, IL-1β, TNF-α, and IL-6^9,11^. Following HIF-1α induction, IKK activity and subsequent NF-κB signaling could be induced by the kinase transforming growth factor-b-activated kinase (TAK1)^12^. Interestingly, NF-κB signaling can promote both OXPHOS and glycolysis depending on the context^13^. For example, p65 (RelA) has been shown to drive increased mitochondrial OXPHOS ^14^, but enhance glycolysis upon loss of p53^15^. Additionally, p53 could play a role in macrophage activation through its effects on oxidative stress^16-19^. By promoting OXPHOS via synthesis of cytochrome c oxidase 2 activation^20^, p53 can boost mitochondrial ROS generation through cytoplasmic polyadenylation element-binding protein^21^. Although elevated glycolytic flux and p53 expression may appear mutually exclusive^22^, p53 could complex with HIF-1α for downstream signaling^23^.

While LPS exposure models involving an initial low dose followed by a high dose over two days offer valuable insights^24,25^, they often fail to fully replicate the clinical characteristics of tolerogenicity, where anti-inflammatory cytokines are concurrently elevated and macrophages exhibit impaired phagocytosis^26,27^. In contrast, models employing long-term LPS exposure (for 7 days) more accurately capture a comprehensive and clinically relevant phenotype of tolerance^26,27^. However, the role of immunometabolism in these models of persistent and chronic inflammation remains undercharacterized, especially when compared to the well-studied immunometabolic changes following short-term (≤ 3 days) LPS exposure^5,6,28,29^. Understanding the immunometabolism of persistent LPS exposure is crucial because this prolonged stimulation of TLR4 agonists is a key driver in various chronic inflammatory conditions, including lung diseases^30-32^ like progressive pulmonary obstructive conditions and chronic bronchitis, chronic inflammation triggered by particle-releasing implanted biomaterials^8,33-35^, inflammatory bowel disease and chronic hepatitis characterized by sustained exposure to gut-derived LPS^36^. Furthermore, the inflammatory profile of macrophages differs significantly between short-term and long-term LPS stimulation, with prolonged exposure often leading to a more tolerogenic phenotype^26,27^. Given the clear clinical relevance of persistent TLR4 agonist exposure and the observed distinct inflammatory responses, elucidating the bioenergetics and functional metabolic profile of persistently activated macrophages, and how these differ from their transiently activated counterparts, as well as the functional significance of NF-κB signaling and p53 activity in these persistent inflammation models, is essential.

Here, we show that persistent LPS stimulation of macrophages, unlike acute exposure, leads to increased oxygen consumption alongside reduced ATP and elevated glycolysis, with a mixed inflammatory profile. Pharmacologically blocking complex I lowered oxygen consumption and diminished mitochondrial ROS, potentially by affecting NF-κB signaling. Unexpectedly, NF-κB inhibitors decreased both oxygen consumption and glycolytic flux with little effect on ROS, while p53 activation similarly reduced oxygen consumption and glycolytic flux and slightly lowered ROS. Ultimately, the lack of elevated mitochondrial membrane potential in prolonged LPS exposure indicated that ROS production was likely not driven by reverse electron transport in this chronic setting, a conclusion supported by the ineffectiveness of a mitochondrial depolarizer on oxygen consumption.

## Results and Discussion

To understand how the immunometabolic signatures of acute (3 days) versus persistent (7 days) exposure of murine bone marrow-derived macrophages (BMDMs) to bacterial lipopolysaccharide (LPS) differ, we characterized ATP levels, functional metabolic profiles, and cell numbers (Fig. 1a-h). Consistent with prior findings^3,5,6^, our results show that acute LPS exposure in BMDMs leads to reduced ATP levels^8^, increased extracellular acidification rate (ECAR, indicative of glycolytic flux), and decreased oxygen consumption rate (OCR, a measure of OXPHOS), changes that are not attributable to alterations in cell numbers (Fig. 1a-d). Extending these observations, we found that persistent LPS exposure in BMDMs similarly results in reduced ATP levels and elevated glycolytic flux, independent of cell number changes (Fig. 1e-g). However, in contrast to acute exposure, persistent stimulation also led to increased oxygen consumption (Fig. 1f), similar to observations made in chronic inflammatory conditions^37-41^. Cytokine and chemokine profiling of persistently-stimulated macrophages showed elevated levels of the pro-inflammatory proteins IL-1β (Fig. 2a) and MCP-1 (Fig. 2b). Notably, TNF-α and IL-6 levels were reduced (Fig. 2c-d), while the expression of the anti-inflammatory cytokines IL-4 and IL-10 was increased (Fig. 2e-f). The mixed inflammatory phenotype is also observed in human macrophages from peripheral blood mononuclear cells and is likely a consequence of endotoxin tolerance^26^. Mechanistically, tolerogenic phenotypes are due to co-expression of anti-inflammatory cytokines^42^ and the down-regulation of IL-6 and TNF-α transcription, RNA polymerase II recruitment and NF-κB and CCAAT/enhancer-binding protein β transcription factor binding to the IL-6 and TNF-α promoters in tolerized immune cells^43^.

**Figure 1:**
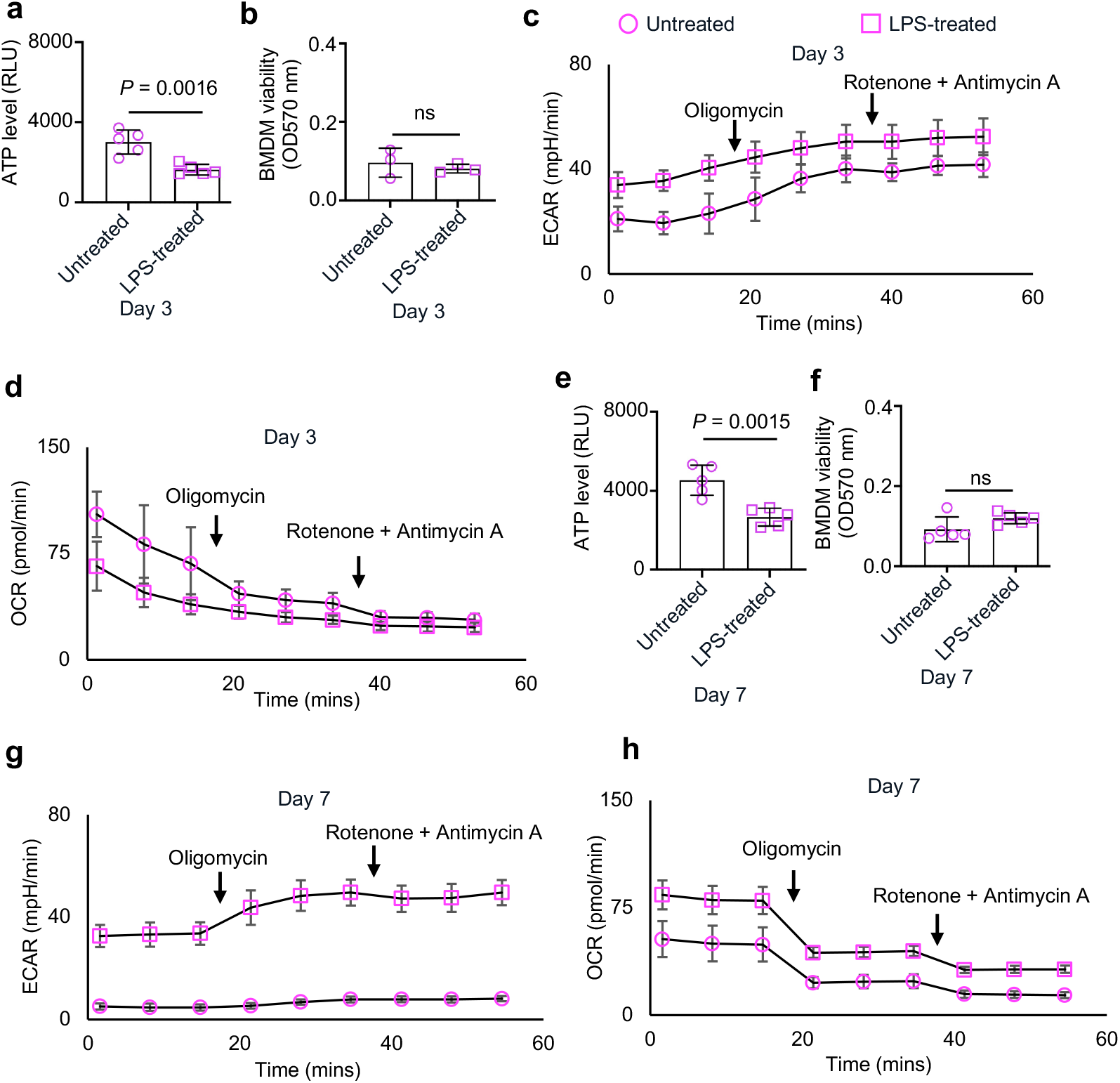
Alterations in bioenergetics and functional metabolic profiles of macrophages with varying endotoxin exposure are not due to changing cell numbers. **a-b**, Adenosine triphosphate (ATP) levels and cell viability of primary bone marrow-derived macrophages (BMDMs) exposed to lipopolysaccharide (LPS) for 3 days. **c-d**, Extracellular acidification rate (ECAR) and oxygen consumption rate (OCR) in BMDMs exposed to LPS for 3 days. **e-f**, ATP levels and BMDM viability following exposure to LPS for 7 days. **g-h**, ECAR and OCR in BMDMs exposed to LPS for 7 days. Mean (SD), n= 3-5 replicates, two-tailed unpaired t-test, not significant (ns).

**Figure 2:**
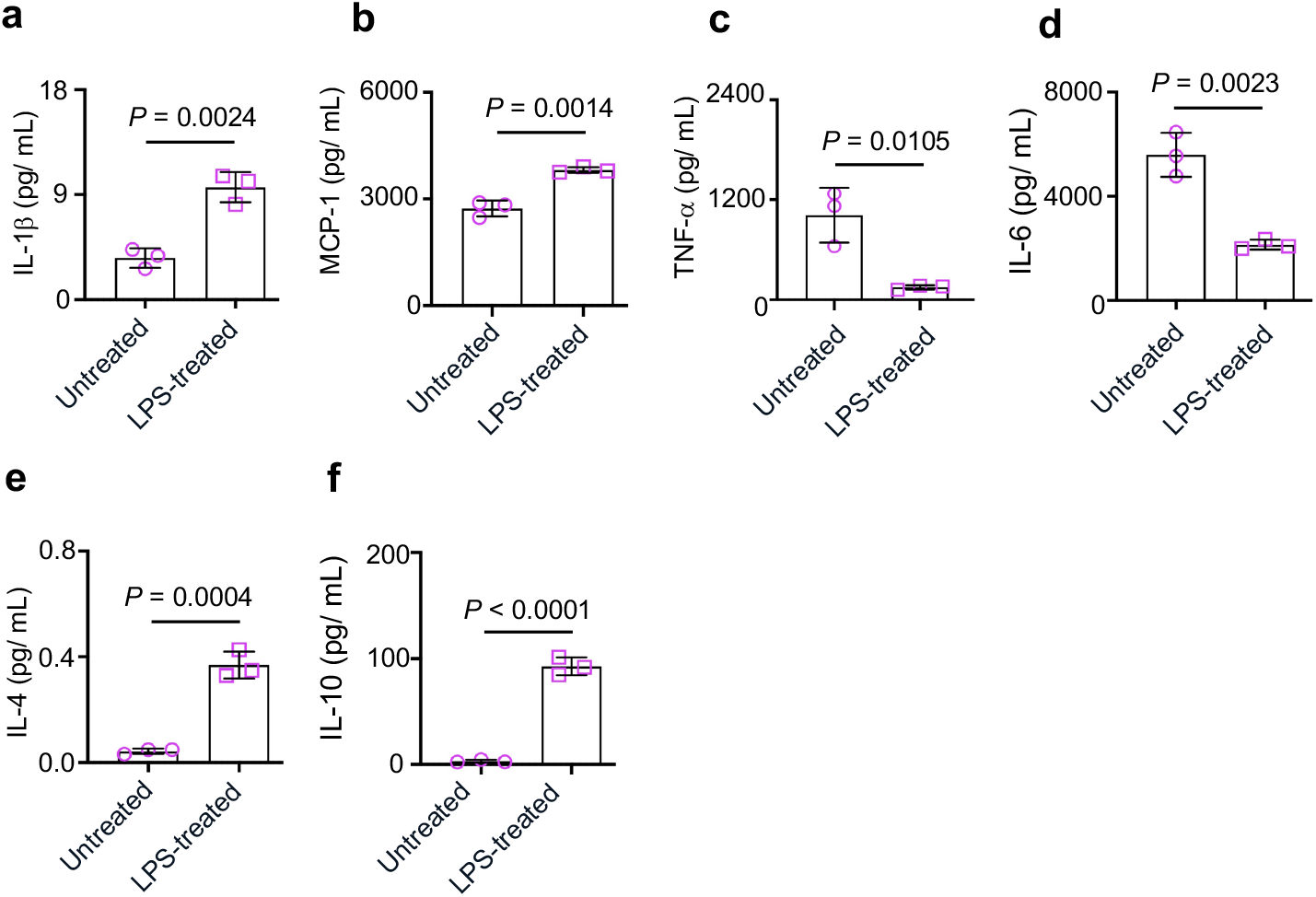
Persistent endotoxin exposure induces simultaneous expression of both pro-inflammatory and anti-inflammatory profiles in BMDMs. **a-f**, IL-1β, MCP-1, TNF-α, IL-6, IL-4 and IL-10 protein levels in macrophages exposed to LPS for 7 days. Mean (SD), n = 3, two-tailed unpaired t-test.

To investigate the functional significance of elevated oxygen consumption in activated BMDMs persistently exposed to LPS, we used several pharmacological agents targeting key pathways. First, given that RET at complex I of the mitochondrial electron transport chain (ETC) is a source of ROS and its inhibition by rotenone reduces oxygen consumption^6^, we used rotenone to assess the contribution of this process. Second, because NF-κB signaling via RelA is known to drive oxygen consumption^14^, we examined the impact of NF-κB inhibition using two distinct agents: BAY 11-7082, which inhibits IκBα phosphorylation essential for NF-κB activation^9,11^, and 5Z-7-oxozeaenol, which blocks TAK1^44,45^, an upstream kinase for IKK activation and subsequent NF-κB signaling^12^. Finally, considering the established role of p53 as a key regulator of OXPHOS^20^, we utilized nutlin-3, a chemical agent that stabilizes and activates p53^46,47^, to determine if enhancing p53 activity would modulate oxygen consumption in our model of persistent inflammation.

We reproduced our previous observation that persistent exposure of BMDMs to LPS simultaneously increased basal OCR and ECAR (Fig. 3a-b). Subsequently, we found that treatment with BAY 11-7082, 5Z-7-oxozeaenol, and nutlin-3 each reduced both OCR and ECAR. While NF-kB signaling is thought to be refractory following prolonged exposure to LPS^48^, in acute LPS exposure, both BAY 11-7082 and 5Z-7-oxozeaenol, via NF-kB inhibition, are anti-inflammatory^9,11,44,45^. Interestingly, by enhancing p53 activation, nutlin-3 downregulates anti-inflammatory gene expression and antagonizes the development of tolerance following prolonged LPS exposure^49^. In contrast, rotenone treatment decreased OCR in a dose-dependent manner without affecting ECAR (Fig. 3a-b). Since oxygen consumption could be related to mitochondrial ROS generation^6,8,50,51^, we next assessed the roles of complex I inhibition, NF-κB inhibition, and p53 stimulation on mitochondrial ROS production in our persistent inflammation model using flow cytometry. Consistent with prior findings in acute LPS exposure^6,8^, persistent LPS exposure elevated mitochondrial ROS levels compared to untreated BMDMs (Fig. 3c-d). However, compared to BMDMs exposed to LPS alone, BAY 11-7082 showed no effect on mitochondrial ROS production, while 5Z-7-oxozeaenol and nutlin-3 mildly reduced it (Fig. 3c-d). Our observed reduction of OCR and mitochondrial ROS by nutlin-3 treatment contrasts observations in normal lymphocytes where p53 activation drives mitochondrial activity and ROS production^52^. Notably, rotenone potently reduced mitochondrial ROS production (Fig. 3c-d). Interestingly, western blot analysis revealed that rotenone reduced both total and phosphorylated p65 levels without affecting p53 levels (Fig. 3e), suggesting that complex I inhibition may reduce mitochondrial ROS by targeting NF-κB signaling.

**Figure 3:**
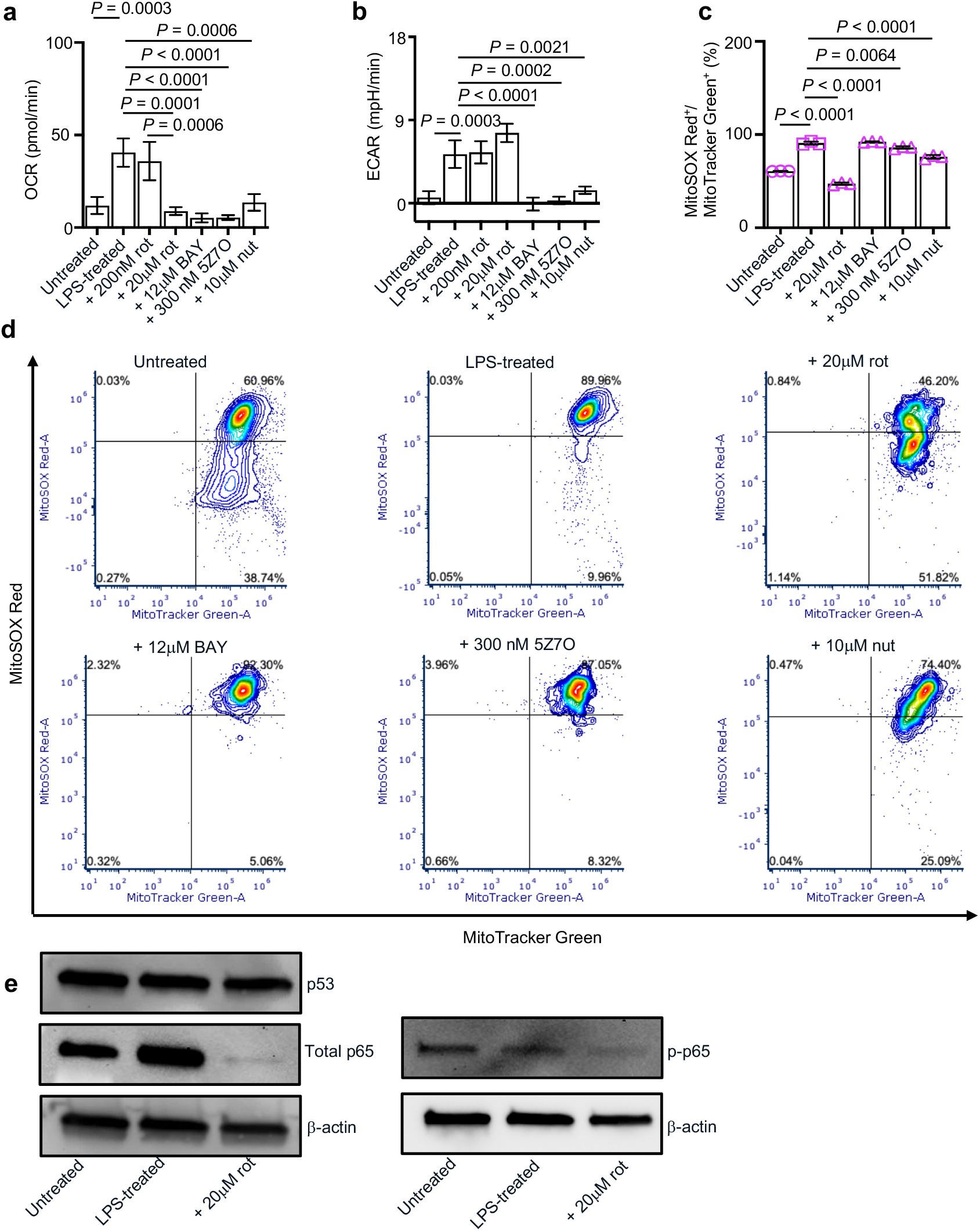
Modulation of oxygen consumption at complex I reduces mitochondrial reactive oxygen species (ROS) production and NF-kB signaling, but not p53 expression in persistently-activated macrophages. **a-b**, Basal oxygen consumption rate (OCR) and extracellular acidification rate (ECAR) of primary bone marrow-derived macrophages (BMDMs) exposed to lipopolysaccharide (LPS), with and without, rotenone (rot), BAY 11-7082 (BAY), (5Z)-7-oxozeaenol (5Z7O) and nutlin-3 (nut) for 7 days. **c-d**, Mitochondrial ROS as measured by MitoSOX Red gated on MitoTracker Green, with representative fluorescence-activated cell sorting plots. **e**, Expression of p53, total p65, phospho-p65 and β-actin (loading control) by western blotting on whole-cell lysates. Mean (SD), n = 3, one-way ANOVA followed by Tukey’s post-hoc test.

Given that mitochondrial hyperpolarization (membrane potential) is a major driver of RET-dependent mitochondrial ROS generation following acute LPS exposure^6-8^, we assessed mitochondrial membrane potential by flow cytometry. We observed that persistent LPS exposure in BMDMs did not lead to increased membrane potential (Fig. 4a-b), suggesting that RET may not be the primary driver of mitochondrial ROS in our persistent inflammation model, an observation previously seen with LPS exposure models involving an initial low dose followed by a high dose^24^. To further evaluate the potential contribution of mitochondrial membrane potential to ROS production, we treated BMDMs persistently exposed to LPS with the mitochondrial depolarizer carbonyl cyanide-4-(trifluoromethoxy)phenylhydrazone^6^. This treatment showed no effects on either OCR or ECAR (Fig. 4c-d), further supporting the idea that increased oxygen consumption is not linked to mitochondrial membrane potential. Interestingly, while rotenone and 5Z-7-oxozeaenol decreased membrane potential, BAY 11-7082 and nutlin-3 increased mitochondrial membrane potential (Fig. 4b), suggesting a nuanced role for NF-κB inhibition at IκBα phosphorylation and TAK1 for dissipating mitochondrial membrane potential in this model of persistent inflammation.

**Figure 4:**
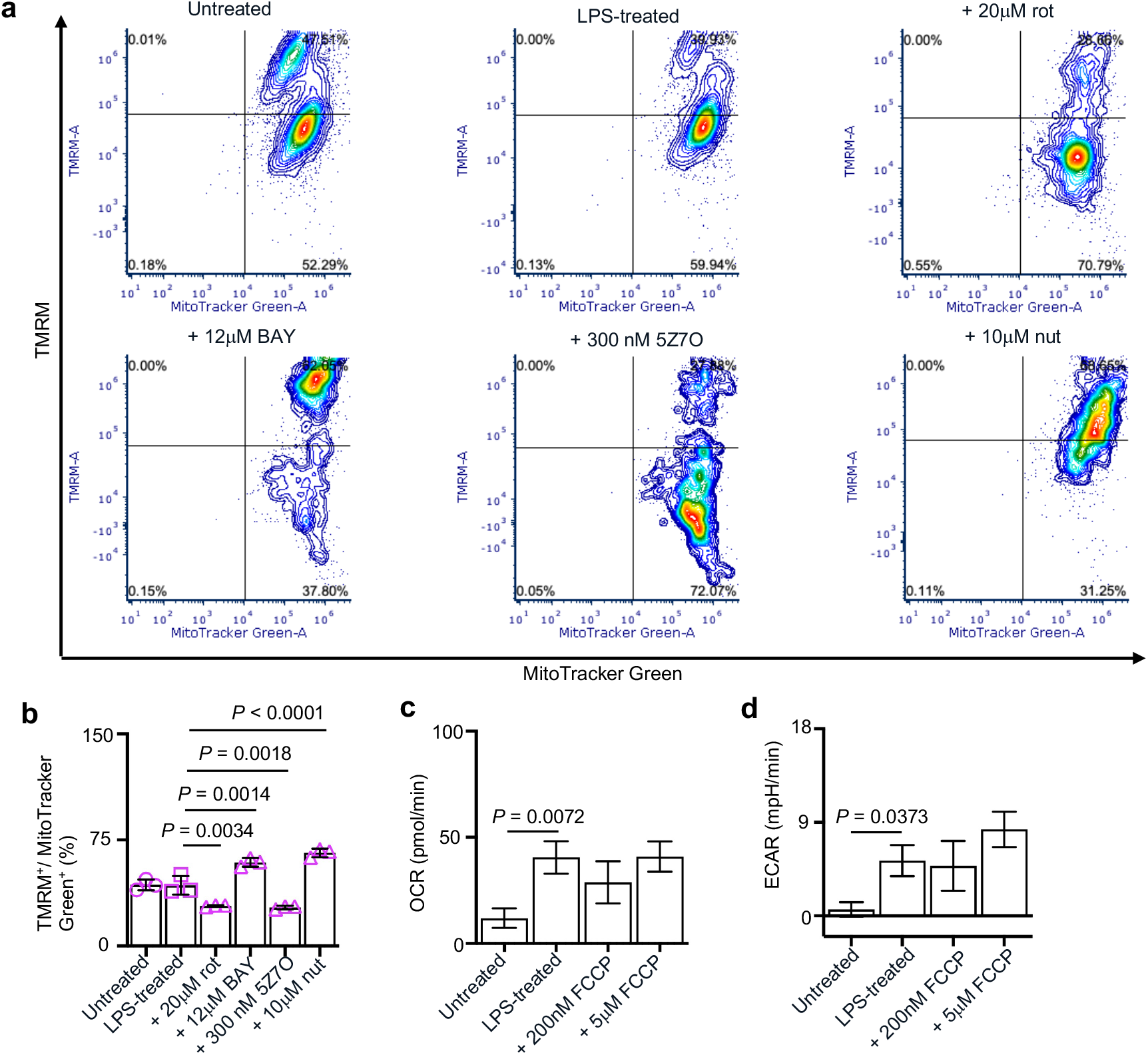
Although their elevated oxygen consumption remains unaffected by mitochondrial uncoupling, persistently activated macrophages display normal mitochondrial membrane potential, which is reduced by complex I inhibition, increased by p53 stimulation and differentially affected by NF-kB inhibition. **a-b**, Mitochondrial membrane potential, quantified by tetramethylrhodamine methyl ester (TMRM) fluorescence gated on MitoTracker Green and shown in representative fluorescence-activated cell sorting plots, was assessed in primary bone marrow-derived macrophages (BMDMs) treated for 7 days with lipopolysaccharide (LPS), alone or in combination with rotenone (rot), BAY 11-7082 (BAY), (5Z)-7-oxozeaenol (5Z7O), or nutlin-3 (nut). **c-d**, Basal oxygen consumption rate (OCR) and extracellular acidification rate (ECAR) of BMDMs exposed to lipopolysaccharide (LPS), with and without, the mitochondrial uncoupler, carbonyl cyanide-p-trifluoromethoxyphenylhydrazone (FCCP). Mean (SD), n = 3, one-way ANOVA followed by Tukey’s post-hoc test.

In summary, we compared the immunometabolic profiles of macrophages under acute (3 days) and persistent (7 days) LPS exposure, revealing that while both showed reduced ATP and increased glycolysis, persistent exposure uniquely exhibited increased oxygen consumption and a mixed inflammatory cytokine profile. Investigating the elevated oxygen consumption in persistent inflammation, pharmacological inhibition of complex I reduced OCR and potently reduced mitochondrial ROS, potentially by impacting NF-κB signaling. Surprisingly, NF-κB inhibitors reduced both OCR and ECAR but had minimal impact on ROS, while p53 activation also reduced OCR and ECAR and mildly reduced ROS, contrasting its typical effects in lymphocytes. Finally, the absence of increased mitochondrial membrane potential in persistent LPS exposure suggested that ROS generation was likely independent of RET in this chronic model, a finding corroborated by the lack of effect of a mitochondrial depolarizer on oxygen consumption rate, while rotenone and 5Z-7-oxozeaenol decreased membrane potential, and BAY 11-7082 and nutlin-3 increased it.

## Methods

### Chemicals

The following chemical inhibitors and activators were used in this study at concentrations noted in each figure legend: rotenone, Bay 11-7082, and (5Z)-7-Oxozeaenol (all from Sigma-Aldrich); carbonyl cyanide-p-trifluoromethoxyphenylhydrazone (Abcam); and nutlin-3 (Santa Cruz Biotechnology).

### Isolation and culture of primary macrophages

Primary bone-marrow derived macrophages (BMDMs) were isolated from male and female C57BL/6J mice (Jackson Laboratories) of 3-4 months as previously described^53^. Isolated (adherent) BMDMs were seeded at 50,000 cells per well of a 96-well plate in complete medium made of DMEM or DMEM F-12 medium, 10% heat-inactivated Fetal Bovine Serum and 100 U/mL penicillin-streptomycin (all from ThermoFisher Scientific) for either 3 or 7 days, as indicated in figure legends. In addition to treating wells with 100 ng/mL of lipopolysaccharide (LPS) from *Escherichia coli* O111:B4 (Sigma-Aldrich) or leaving them untreated as controls, specific wells received chemical inhibitors or activators dissolved in dimethyl sulfoxide (DMSO; Sigma-Aldrich). To ensure consistency, untreated control wells also contained DMSO. The final concentration of each chemical within the wells is specified in the corresponding figure legends.

### ATP assessment

Macrophage ATP levels were quantified using commercially available assay ATP/ADP kits (Sigma-Aldrich). These kits, containing D-luciferin, luciferase, and a cell lysis solution, were utilized following the manufacturer’s protocol. Luminescence was measured with an integration time of 1000 ms using a SpectraMax M3 Spectrophotometer (Molecular Devices) and analyzed with SoftMax Pro software (Version 7.0.2, Molecular Devices).

### Seahorse assays

Real-time measurements of basal oxygen consumption rate (OCR) and extracellular acidification rate (ECAR) were undertaken using either the Seahorse XFp or XFe-96 Extracellular Flux Analyzer (Agilent Technologies. Before the assay, cell culture medium was removed and cells were washed and incubated in Seahorse XF DMEM medium (pH 7.4) reconstituted to contain 25 mM D-glucose and 4 mM Glutamine. Seahorse plates were equilibrated in a non-CO_2_ incubator for one hour prior to measurement. The Seahorse ATP rate assay were performed following the manufacturer’s guidelines and final well concentrations of 1.5μM and 0.5μM of oligomycin and rotenone/antimycin A, respectively, were sequentially injected after obtaining baseline measurements. All reagents for the Seahorse assays were obtained from Agilent Technologies. Seahorse data, expressed as means ± standard deviation (SD), were exported directly using Wave software (Version 2.6.1).

### Macrophage viability

Cell viability was determined at the end of the culture period by quantifying adherent cell biomass using the crystal violet staining assay^54^ performed at ambient temperature. In brief, 150 µl of the 200 µl medium in each well of a 96-well plate was removed. Cells were then fixed and permeabilized by adding 150 µl of 99.9% methanol (MilliporeSigma) for 15 minutes, which was subsequently removed. Following this, cells were stained with 100 µl of a 0.5% crystal violet solution in 25% methanol for 20 minutes, after which the wells were washed. Each well underwent two 2-minute washes with 200 µl of phosphate buffered saline. The resulting absorbance, or optical density, was measured at 570 nm using a SpectraMax M3 Spectrophotometer (Molecular Devices) and analyzed with SoftMax Pro software (Version 7.0.2, Molecular Devices).

### Cytokine and chemokine profiling

Supernatant concentrations of cytokines and chemokines were pooled then quantified using a MILLIPLEX MAP mouse magnetic bead-based multiplex assay kit (MilliporeSigma) in triplicate^55^. This kit was employed to determine the protein expression levels of IL-6, MCP-1, TNF-α, IL-1β, IL-4, and IL-10. Data acquisition was performed using a Luminex 200 instrument (Luminex Corporation) controlled by xPONENT software (Version 3.1, Luminex Corporation).

### Flow cytometry

Following incubation using 1X PBS containing 4mM EDTA (Teknova) at 37 °C for 10 minutes, macrophages were detached from 96-well plates using a cell scraper aided by careful pipetting. The cells were pooled before analyzing analyzed for mitochondrial mass (MitoTracker Green FM, MTG; ThermoFisher Scientific), mitochondrial transmembrane potential (tetramethylrhodamine methyl ester, TMRM; ThermoFisher Scientific), and mitochondrial superoxide levels (MitoSOX Red, MSOX; ThermoFisher Scientific). Control samples included unstained cells for each condition and cells stained with individual dyes (TMRM, MTG, or MSOX). For staining, 650,000 cells in 100 µL of 1X PBS (ThermoFisher Scientific) containing Live/Dead Blue (1:1000) and either MTG (50 nM)/TMRM (20 nM) or MTG (50 nM)/MSOX (2.5 uM) were incubated in the dark at 37 °C with 5% CO2 for 30 minutes. Cells were pelleted by centrifugation and washed twice with flow buffer (0.5% bovine serum albumin in 1X PBS; MilliporeSigma). Subsequently, cells were fixed with 4% paraformaldehyde for 10 minutes at room temperature in the dark, followed by centrifugation and resuspension in 100 µL of flow buffer for analysis in triplicate using a Cytek Aurora flow cytometer. Cell populations were identified and single cells selected using forward and side scatter. MTG-positive cells were gated from the live cell population, and MSOX/TMRM signals were assessed within the MTG-positive subset. Data analysis was performed using FCS Express software (De Novo Software; version 7.12.0005).

### Western blotting

Mouse BMDMs were collected, pooled and lysed using RIPA lysis buffer (ThermoFisher Scientific). Protein levels in the lysates were quantified using the Pierce BCA Protein Assay kit (ThermoFisher Scientific) following the provided protocol. Equal amounts of protein were mixed with 4X Laemmli sample buffer (Bio-Rad) and heated at 95°C for 5 minutes. The denatured protein samples were then separated by electrophoresis on 4-20% SDS-PAGE gels (Bio-Rad) and subsequently transferred to 0.2 μm polyvinylidene difluoride (PVDF) membranes using a Turbo semi-dry transfer system (Bio-Rad). To minimize non-specific antibody binding, membranes were blocked with Everyblot Blocking Buffer (Bio-Rad) for a minimum of 10 minutes at room temperature, followed by an overnight incubation at 4°C with the indicated primary antibodies: rabbit monoclonal anti-NF-κB p65 (D14E12) XP #8242 (Cell Signaling Technology); rabbit monoclonal anti-phospho-NF-κB p65 (Ser536) (93H1) #3033 (Cell Signaling Technology); mouse monoclonal anti-p53 (1C12) #2524 (Cell Signaling Technology); and mouse monoclonal anti-β-Actin # A1978 (Sigma-Aldrich). After several washes with 0.1% Tris-buffered saline with Tween-20 (TBST), membranes were incubated with appropriate horseradish peroxidase (HRP)-conjugated secondary antibodies. Following further washes, protein bands were detected using an enhanced chemiluminescent (ECL) substrate (Bio-Rad) and visualized using a ChemiDoc MP imaging system (Bio-Rad). Three independent western blot measurements were undertaken.

### Statistics

Data, presented as the mean with standard deviation (SD), were analyzed using GraphPad Prism software (Version 9.5.1 (528)). Specific statistical tests employed, resulting p-values, and the number of samples in each group are detailed within the corresponding figure legends.

## Data availability

The data supporting the findings of this study are available within the paper.

## Acknowledgements

Funding for this work was provided in part by the James and Kathleen Cornelius Endowment at MSU. Flow cytometry was performed in the MSU Cytometry Core.

## Author contributions

Conceptualization, C.V.M. and C.H.C.; Methodology, C.V.M., X.X., A.V.M., E.U., and C.H.C.; Investigation, C.V.M., X.X., A.V.M., and E.U.; Writing – Original Draft, C.V.M.; Writing – Review & Editing, C.V.M., X.X., A.V.M., E.U., and C.H.C.; Funding Acquisition, C.H.C.; Resources, C.H.C.; Supervision, C.H.C.

